# Mainly on the Plane: Deep Subsurface Bacterial Proteins Bind and Alter Clathrate Structure

**DOI:** 10.1101/2020.06.15.152181

**Authors:** Abigail M. Johnson, Dustin J.E. Huard, Jongchan Kim, Priyam Raut, Sheng Dai, Raquel L. Lieberman, Jennifer B. Glass

## Abstract

Gas clathrates are both a resource and a hindrance. They store massive quantities of natural gas but also can clog natural gas pipelines, with disastrous consequences. Eco-friendly technologies for controlling and modulating gas clathrate growth are needed. Type I Antifreeze Proteins (AFPs) from cold-water fish have been shown to bind to gas clathrates via repeating motifs of threonine and alanine. We tested whether proteins encoded in the genomes of bacteria native to natural gas clathrates bind to and alter clathrate morphology. We identified putative clathrate-binding proteins (CBPs) with multiple threonine/alanine motifs in a putative operon (*cbp*) in metagenomes from natural clathrate deposits. We recombinantly expressed and purified five CbpA proteins, four of which were stable, and experimentally confirmed that CbpAs bound to tetrahydrofuran (THF) clathrate, a low-pressure analog for structure II gas clathrate. When grown in the presence of CbpAs, THF clathrate was polycrystalline and plate-like instead of forming single, octahedral crystals. Two CbpAs yielded branching clathrate crystals, similar to the effect of Type I AFP, while the other two produced hexagonal crystals parallel to the [1 1 1] plane, suggesting two distinct binding modes. Bacterial CBPs may find future utility in industry, such as maintaining a plate-like structure during gas clathrate transportation.

**Table of Contents Graphic:** 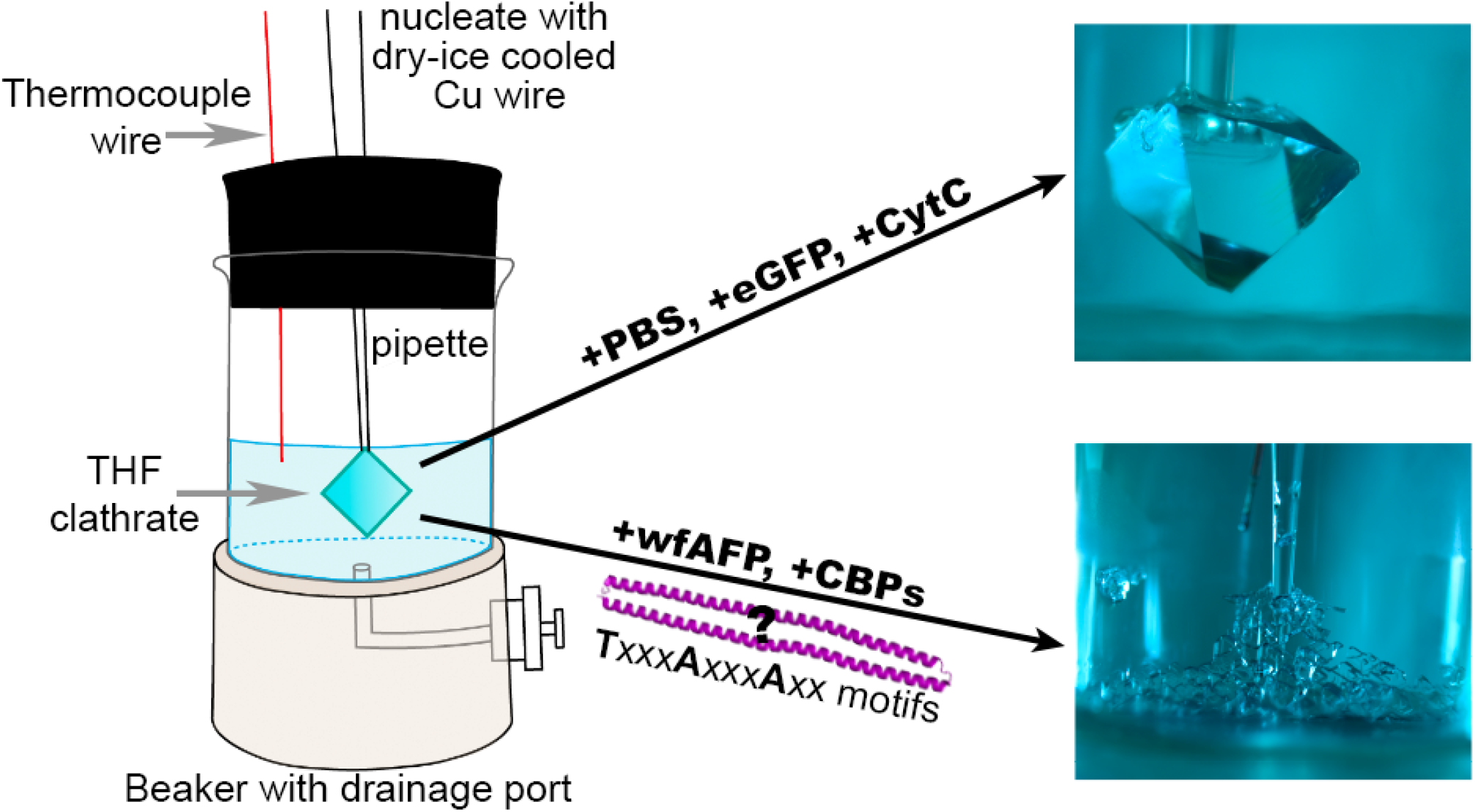

---

Gas clathrates—crystalline structures of hydrogen-bonded water molecules that encage various gases via van der Waals□ interactions—are found along continental margins, in and under permafrost, and likely on a number of other planetary bodies throughout the solar system^1-3^. Gas clathrates have garnered considerable interest for their implications in climate change^4-6^ and as prospective energy resources^7^. The natural gas industry devotes considerable financial resources^8^ to synthetic chemical inhibitors^9^ of gas clathrate formation because clogged natural gas pipelines pose human and environmental safety^10, 11^ hazards (e.g. the Deepwater Horizon Oil Spill).

In the search for more-“green” gas clathrate inhibitors, antifreeze proteins (AFPs) were found to provide superior clathrate inhibition than synthetic commercial inhibitors^12^. AFPs enable diverse organisms, from bacteria to fish, to survive under low-temperature conditions by binding specific ice planes irreversibly, thereby depressing the freezing point of ice^13-16^. Although ice and gas clathrates have different crystalline structures, Type I AFPs inhibit gas clathrate^12, 17-22^. Type I AFPs bind clathrates using the motif TxxxAxxxAxx, where x is any amino acid^23^.

Gas clathrates are known to support microbial life^24^. Gas clathrate-dwelling archaea and bacteria were found to be physically associated with the gas clathrates at average concentrations of 10^6^ cells mL^-1 24^. Here we report the first characterization of clathrate binding proteins (CBPs) encoded in bacterial genomes from gas clathrate stability zones.

We identified five potential *cbp* genes (*cbpA_2,3,5,6,8_*) within conserved gene clusters (*cbpBCD(A□,A□ □)A*, **Figure 1A; Table S1**) from metagenome analysis of Hydrate Ridge (ODP site 1244), offshore Oregon^25^, and other gas clathrate-rich sites including offshore Shimokita Peninsula, Japan^26^. Gene products of *cbpA* share secondary structure similarity with Maxi, the larger, hyperactive isoform of the winter flounder *(Pseudopleuronectes americanus)* AFP^27^. CbpAs harbor one to six Type I AFP clathrate binding motifs, particularly in their conserved C-terminal domains, which are predicted^28^ to form coiled-coils (**Figure 1B, C**). CbpAs have conserved N- and C-termini (orange and green, **Figure 1A, C**), though the former is absent in CbpA_8_ and found elsewhere (A□/A□ □) in the contigs of CbpA_3,5_. All CbpAs except CbpA_2_ share a common N-terminal domain (purple, **Figure 1A, C**), but none harbor secretion signal sequences that are readily detected by prediction software^29^. Reminiscent of other Type I AFPs,^30^ CbpAs are largely composed of alanine residues (32.8-42.4% composition, **Table S2**) and are rich in prolines (up to 5.6%, **Table S2**). Pro-rich areas cluster at the end of the N-terminal domain (CbpA_3,5,6,8_), or near the end of the C-terminal domain (magenta, **Figure 1A, C**).

**Figure 1.**
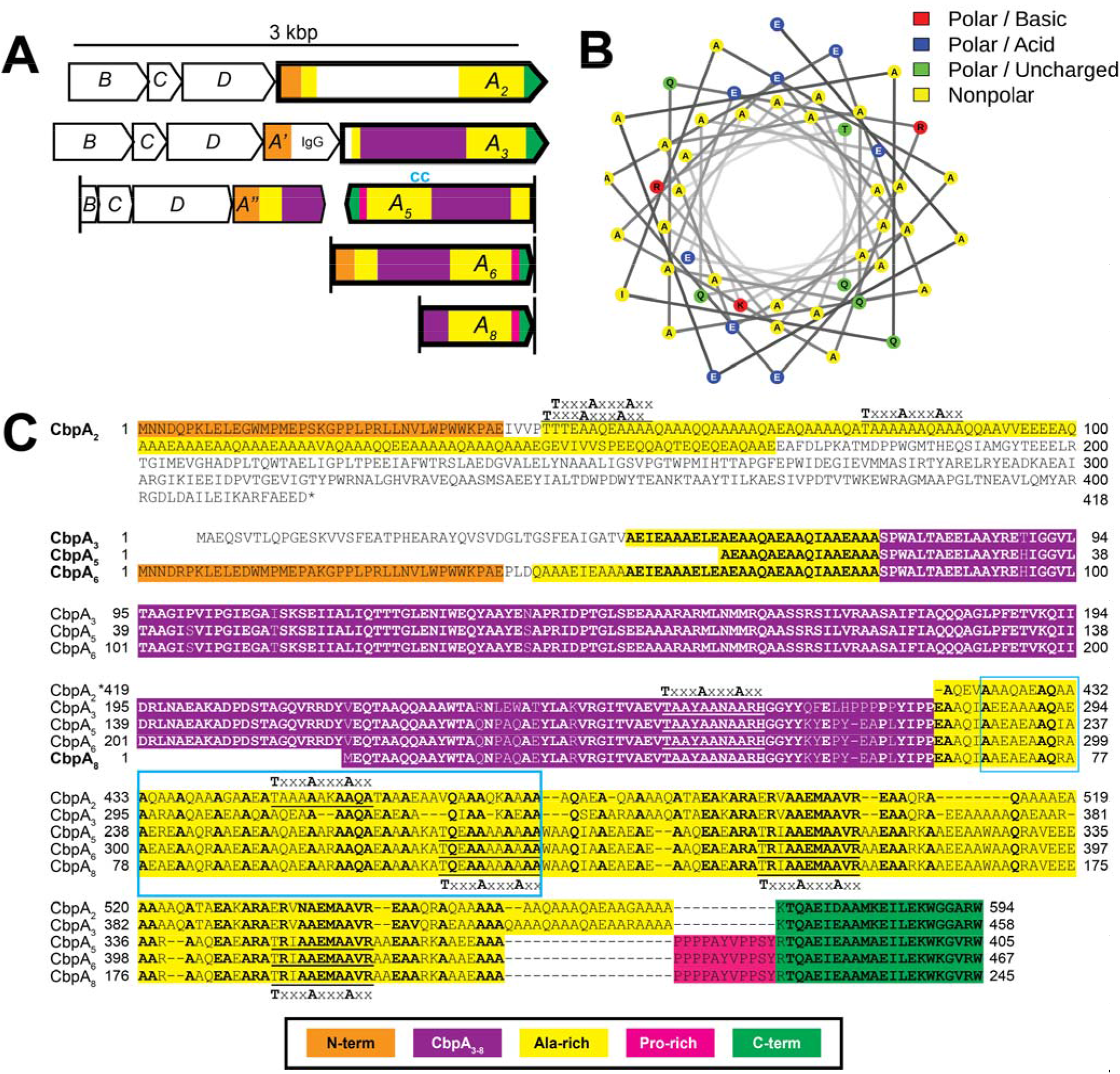
Gene arrangement and CbpA alignment. A) *cbpBCDA* gene clusters from five metagenomes from methane clathrate-bearing sediments. Genes encoding CbpA proteins that were the focus of this study are marked with bold borders and labeled A_2_, A_3_, A_5_, A_6_, and A_8_. *cbpA_3_’* and *cbpA_5_”* denote genes upstream from *cbpA.* The “cc” label in *cbpA*_5_ depicts the best predicted coiled coil, shown in (B). B) Coiled coil region, labeled by type of amino acid. C) Amino acid alignments of the five CbpA proteins. Colored regions correspond to the color scheme in (A). The blue box in CbpA5 denotes the region of the helical wheel in (B). Type I AFP motifs (TxxxAxxxAxx) are underlined in CbpA sequences.

While the importance and interrelatedness of the gene products in these putative *cbp* operons remain unclear, neighboring genes encode an apparent cysteine peptidase (*cbpB)* and a bacterial cell adhesion protein (**Tables S3, S4**). The *cbp* gene sequences most likely originated from *Dehalococcoidia* bacteria, of the phylum *Chloroflexi* (**Table S3, S4**). *Chloroflexi*, which have gained interest for their capacity to metabolize organohalides^31-33^, are known to populate methane clathrate-rich sediment^26, 34, 35^.

We expressed and purified the putative CbpAs to homogeneity (**Figure S1**). CpbA_8_, the shortest gene product (**Figure 1A, C**), degrades appreciably within 24 h (data not shown), and was not studied further. CbpA_2,3,5,6_ exhibit strong α-helical secondary structure signatures by circular dichroism (**Figure S1**) and migrate in SDS-PAGE as species larger than their expected molecular masses (**Figure S1**, **Table S5**). Finally, CbpAs exhibit thermal melting temperatures below 40°C (**Figure S1, Table S6**).

Next, we tested for gas clathrate activity using an established surrogate assay using liquid THF clathrate. Although THF clathrate does not form in nature, it is commonly used as a gas clathrate analog due to its low-pressure stability to study potential natural gas clathrate inhibitors^17,18,36-42^. THF clathrate adopts structure II (sII) clathrate with octahedral morphology and is composed of two types of cavities - 16 small pentagonal dodecahedron (5^12^) and 8 large pentagonal hexadecahedron (5^12^6^4^) cages^10^. We grew THF clathrate at −3°C from a stoichiometric solution^43^ composed of 19.1 weight percent (wt%) THF in the presence or absence of additives, using the setup presented in **Figure S2**. To probe for additive bound to clathrate, concentrations in the solution remaining after crystal growth and in the melted clathrate were measured at the end of each experiment. Negative controls with no additives (**Figure S3A**), phosphate-buffered saline solution, cytochrome c, and eGFP treatments resulted in single octahedral crystals, or twinned crystals with clearly defined [1 1 1] faces **(Figure 2A)**. Cytochrome c and eGFP treatments resulted in more protein in solution compared to clathrate; the average [protein] in clathrate melt: [protein] in non-crystallized solution (hereafter [C/S]) was less than 1 (<0.7; **Figure S4**). Positive controls, including the chemical inhibitor polyvinylpyrrolidone (PVP)^38^ (**Figure S3B**), and Type I AFP **(Figure 2A)** produced, as expected,^17, 18, 36^ twinned or polycrystalline THF clathrate with skeletal or branched morphology, respectively. The [C/S] for Type I AFP was 1.9±0.8, confirming preferential binding.

**Figure 2.**
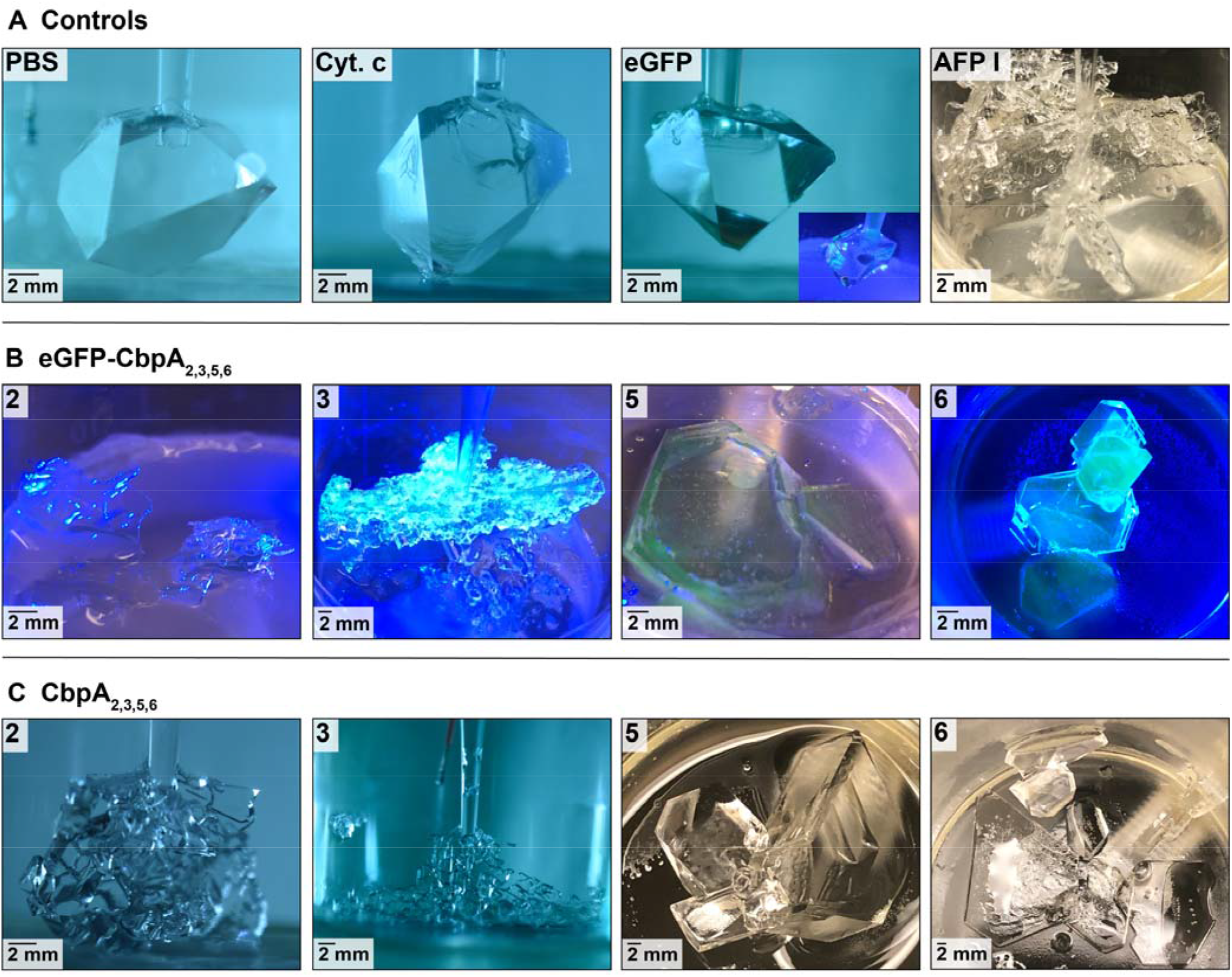
CbpAs induce polycrystalline and plate-like THF clathrate growth. Representative images of each treatment are shown. Protein concentration was 100 μg mL^-1^ for all treatments shown except for eGFP-CbpA_2,5_ (48 and 64 μg mL^-1^, respectively). A) Negative controls: Phosphate buffered saline (PBS) solution, cytochrome c, and eGFP treatments. Positive control: Type I AFP. B) THF clathrate grown in the presence of eGFP-CbpA_2,3,5,6_, with a blue light to illuminate clathrate-bound eGFP. C) THF clathrate grown in the presence of CbpA_2,3,5,6_. Each image is labeled with the treatment or CbpA number and a scale bar.

In general, the addition of CbpAs (CbpA_2,3,5,6_) yielded plate-like morphology of THF clathrate when tested at 100 μg mL^-1^ with or without fusion to eGFP^36, 40-42^ (**Figure 2B, C**), distinct from negative controls (**Figure 2A**, **Figure S3A**)^39^. Under blue light, THF clathrate formed in eGFP-CbpA treatments fluoresced green (**Figure 2B**), indicating the presence of fusion protein bound to the clathrate. Binding was confirmed by [C/S] analysis, which for CbpAs were intermediate between the negative controls and Type I AFP (**Figure S4**). Clathrate crystals grown in the presence of CbpA_2_ or CbpA_3_ were most similar to Type 1 AFP, forming many branched clathrate crystals whereas CbpA_5_ or CbpA_6_ yielded large, flat hexagonal crystals **(Figure 2B, C)**. Concentration dependence was tested for eGFP-CbpA_2_ and CbpA_2_ **(Figure S5)**, revealing a flat plate-like morphology at the higher concentrations (0.53 μM for eGFP-CbpA_2_ and 1.63 μM and 1.22 μM for CbpA_2_) and single THF clathrate crystals with surface deformities, such as kinking, at lower CbpA concentrations (0.35 μM for eGFP-CbpA_2_ and 0.82 μM for CbpA_2_).

Unexpectedly, morphological changes conferred by CbpAs on THF clathrate structures were dependent on the specific CbpA tested, even though the proteins exhibit high levels of sequence similarity. CbpA_2_ and ChpA_3_ induce branching THF clathrate (**Figure 2B, C**) similar to the effect of the kinetic clathrate inhibitor polyvinylcaprolactam (PVCap) on ethylene oxide clathrate^38^. CbpA_5_ and CbpA_6_, promote 2D hexagonal clathrate growth parallel to the [1 1 1] face **(Figure S6)**, similar to the effect of the winter flounder AFP on THF clathrate^17, 18^.

The differing THF clathrate morphologies obtained in the presence of CbpA_2_ and CbpA_3_ versus CbpA_5_ and CbpA_6_, raise the possibility of two discrete binding modes, such as binding to a different crystalline face, or distinct mechanisms of inhibition. In ice-based systems, AFPs are thought to interact directly with water ice by an absorption inhibition mechanism which involves AFP binding to an actively growing ice front; this confines the space in which additional water can be incorporated into the growing ice lattice, eventually resulting in energetically unfavorable addition and halted ice crystal growth^44,45^. The anchored-clathrate water hypothesis describes a potential avenue of AFP absorption to ice, whereby AFPs organize water molecules into clathrate-like cages that then merge with ice at the ice/fluid water interface^46^; the crystal structure of the dimeric, Type I Maxi AFP supports this method of interaction^27^. Modeling studies predict that simple Type I AFPs bind to clathrate via a similar absorption mechanism, where small hydrophobic side-chains of the AFPs (methyl groups of alanine and threonine from TxxxAxxxAxx motif) fill partially-completed clathrate cages^23^. While our CbpAs have similarities with Maxi their differing effects on THF clathrate morphology should prompt future mechanistic studies into CbpA mode(s) of action, which could shed light on AFP inhibition more generally.

Future studies will extend our protein characterization to other antifreeze properties and our initial focus on the accessible sII THF clathrate. CbpA binding likely extends across the most common gas clathrates, including methane clathrate, which adopts structure I (two small pentagonal dodecahedron (5^12^) and six large hexagonal truncated trapezohedron (5^12^6^2^) cages^10^. First, CbpAs are derived from natural habitats containing methane clathrate. Indeed, higher potency binding may be expected for gas clathrate *in situ*^47^. Second, Type I AFPs found in cold-adapted fish bind can alter both sI^48^ and sII^17, 18^ even though their native substrate is ice. Third, the commercially used clathrate inhibitor PVCap inhibits ethylene oxide clathrate (sI clathrate)^38^, resulting in a similar morphology to those seen for CbpA_2_ and CbpA_3_.

Commercial gas clathrate inhibitors, including the thermodynamic inhibitor methanol, and the kinetic inhibitors PVP and PVPCap, are neither economical nor ecologically friendly^12^. Protein-based alternatives are attractive due to their superior inhibition, and environmentally friendly, biodegradable nature. Although AFPs have been screened for gas clathrate inhibition, AFPs have yet to be commercialized because they cannot be produced in sufficient quantities^12^. Our newly discovered CbpAs isolated from microbial genomes in gas clathrate stability zones that are readily expressed and purified in mg quantities in *E. coli* with likely tailored function to inhibit natural gas clathrates, are an attractive new direction both for gas production from clathrate deposits, as well as controlling crystal growth for gas storage and transport, or for use as a novel separation technique to isolate gases from other hazardous substances, or even as a desalination process^49^. These newly discovered CBPs also have implications for searching for life on other planetary bodies, such as Pluto and Mars, which are predicted to host gas clathrates^3^.

## ASSOCIATED CONTENT

### Supporting Information

The Supporting Information is available free of charge on the ACS Publications website. Supporting information includes protein synthesis methods, experimental procedures, protein biochemical characteristics, protein binding to clathrate, negative controls (PDF).

## ACKNOWLEDGMENT

We thank William Waite (USGS) and Ran Drori (Yeshiva University) for helpful suggestions.

## ABBREVIATIONS

Cbp: clathrate-binding protein
eGFP: enhanced green fluorescent protein
THF: tetrahydrofuran

