## Supplemental Information for "Mainly on the Plane: Deep Subsurface Bacterial Proteins Bind and Alter Clathrate Structure"

**This PDF file includes:**

Materials and Methods

Table S1 to S6

Figures S1 to S6

References 1-28

**Materials and Methods**

**Metagenome mining.** Metagenomes from Hydrate Ridge (ODP site 1244, 68.55 mbsf; BioProject PRJNA390944), sequenced and assembled in another study^1^, was annotated using Prokka^2^ and mined for candidate clathrate-binding proteins (Cbp) by targeting sequences that were similar to that of known AFPs. Sequences similar to Type I AFPs were targeted because they are the most effective AFPs to inhibit gas clathrate formation and growth rate.^3-6^ The pendant methyl groups of the threonine and alanine in the Type I AFP clathrate-binding motif (TxxxAxxxAxx) are predicted to bind the half empty cages of the gas clathrate.^7^ Translated metagenome sequences were searched for every occurrence of “TxxxAxxxAxx” using the match function in the regular expression module for Python v3.7^8^. Sequences with three or more TxxxAxxxAxx motifs were analyzed with AFPredictor^9^ and TargetFreeze^10^ to predict their antifreeze properties based on their predicted structure and primary amino acid sequence, respectively. To obtain additional sequences from other global locations, BLAST in UniProt^11^ was used to search for similar sequences. Sequences with TargetFreeze scores > 0.7 were chosen for overexpression and characterization (see next section). When contigs were of sufficient length, genes upstream from the sequence of interest were analyzed by BLAST to assess taxonomic assignment and by InterProt^12^, HMMER^13^ searching UniProtKB/Swiss-Prot, and Phyre2^14^ for functional assignment. Upstream sequences were concatenated and aligned with MUSCLE using MEGA7^15^. Genes on this putative contig were named *cbp* for clathrate binding protein. The gene encoding the TxxxAxxxAxx motif(s) was designated *cbpA.* Upstream genes were labeled *cbpBCD.* Secondary structure prediction of CbpAs was performed with Jpred 4^16^.

**Recombinant protein expression in *E. coli* and purification.** CbpA proteins from five metagenomes (hereafter CbpA_2,3,5,6,8_) were expressed with an N-terminal hexahistidine tag, followed by thrombin cleavage site, enhanced green fluorescent protein (eGFP)^17^, and tobacco etch virus (TEV) protease cleavage site. Codon-optimized genes coding for CbpA were synthesized and cloned by GenScript into the pET28a(+) vector (Novagen) modified to encode the N-terminal cleavable eGFP fusion. CbpAs were expressed in *E. coli* BL21(DE3) cells cultured in Luria-Bertani broth (BD) supplemented with 50 mgL^-1^ kanamycin sulfate (Sigma-Aldrich). Cells were grown at 37°C and 225 rpm shaking until OD_600_ = 0.6-0.8 was reached, at which point the temperature of the incubator was dropped to 18°C for up to 2 hours. After equilibration at 18°C, cells were induced with 0.5 mM isopropyl-β-D-1-thiogalactopyranoside (IPTG, GoldBio), and were allowed to continue shaking post-induction overnight (~16 hours) prior to harvest via centrifugation (10 min spins at 4,420 x g, 4°C). Pellets were flash-cooled in liquid nitrogen for storage.

eGFP-CbpA_2,3,5,6,8_-containing cell pastes were thawed on ice and resuspended in wash buffer (50 mM Tris, 0.5 M NaCl at pH 8.0) supplemented with protease inhibitor cocktail (cOmplete Tablets EDTA-free, Roche) and deoxyribonuclease I (DNase I, Sigma-Aldrich). Cells were lysed in turn by double passage through a French press. Lysate was then clarified by ultracentrifugation (1 hour at 110,000 x g and 4°C). eGFP-CbpAs were isolated first with a 5 mL HisTrap HP column (GE Healthcare) using an AKTA purification system (GE Healthcare). The column was equilibrated with wash buffer supplemented with 40 mM imidazole (Acros Organics) and eGFP-CbpAs were eluted with a gradient of 40-500 mM imidazole. eGFP-CbpAs bound weakly to the HisTrap column. Next, eGFP-CbpAs were polished by size exclusion chromatography (SEC). eGFP-CbpA_2_ was fractionated on a HiLoad 16/600 Superdex 200 preparatory grade (PG) column (GE Healthcare) whereas eGFP-CbpA_3,5,6,8_ fusions were fractionated on a HiLoad 16/60 Superdex 75 PG column (GE Healthcare). Both columns were equilibrated with wash buffer.

For eGFP-CbpA_3,5,6,8_, eGFP was removed from SEC-purified eGFP-CbpA_3,5,6,8_ fusions by treatment with 1:10 (w/w) TEV protease (prepared in-house using the pRK793 plasmid and published methods^18^) at room temperature for ~16 hours. CbpAs were then isolated by applying the cleavage reaction mixture to the 5 mL HisTrap HP column equilibrated with wash buffer (no imidazole). The His-tagged eGFP and TEV protease bound the column and eluted after applying a 0-500 mM imidazole gradient. eGFP was saved for use in control experiments. CbpA proteins were then purified by SEC as for eGFP-CbpA_3,5,6,8_ fusions.

To purify CbpA_2_, eGFP-CbpA_2_ was dialyzed after HisTrap against 2 L of wash buffer at 4°C for ~16 hours and the first SEC step was omitted. eGFP was then cleaved by treatment with TEV protease, isolated, and eluted as described above. CbpA_2_ was polished by SEC using HiLoad 16/600 Superdex 200 PG equilibrated with wash buffer followed by a final passage over HisTrap HP again equilibrated with wash buffer, to remove all traces of His-tagged contaminants.

The purity of each protein was confirmed by SDS PAGE analysis visualized with Coomassie blue staining. The concentration of eGFP and eGFP-CbpA fusions was estimated by absorbance at 491 nm and published molar extinction coefficient^19^ of 55,000 M^-1^ cm^-1^. Concentrations of CbpAs were determined using the absorbance at 280 nm, using extinction coefficients and molecular weights were obtained from Expasy^20^ (see **Table S1**).

**Secondary structure analysis.** Far-UV circular dichroism (CD) spectra were acquired on a Jasco J-815 spectropolarimeter equipped with a Jasco PTC-4245/15 temperature control system. Cleaved CbpAs were prepared at a concentration range of 0.5-2.0 μM in 10 mM sodium phosphate buffer at pH 7.2 at 4°C. CD scans were measured from 200 nm to 300 nm at a rate of 500 nm/min and a data pitch of 1 nm using a 0.1 cm cuvette. Each measurement was blank subtracted with a buffer control and reflects an average of 10 scans. *In silico* coiled-coil predictions were performed with Waggawagga^21^ servers. Helical wheels were drawn using NetWheels^22^. Signal sequences were predicted using SignalP 5.0^23^.

**Thermal stability measurements.** CbpA thermal stabilities were determined by differential scanning fluorimetry (DSF) using a previous protocol^24^. Briefly, protein stocks were prepared in 10 mM HEPES buffer with 0.2 M NaCl at pH 7.2. Reaction mixtures were prepared as master mixes containing 5x Sypro Orange and a final protein concentration of 1-10 μM in the HEPES buffer. Reaction samples were aliquoted from master mixes (30 μL/well) into a 96-well optical plate (Applied Biosystems) and sealed with optical film. All steps involving Sypro Orange were performed in darkness. Fluorescence measurements were obtained on an Applied Biosciences Step-One Plus real time (RT) PCR instrument. Thermal melts were conducted from 25°C to 95°C with a 1°C temperature increase per every 75 seconds. Data were analyzed using the Boltzmann sigmoid equation in GraphPad Prism 8. The melting temperature (T_m_) was calculated as the midpoint of unfolding. Reported T_m_ values reflect an average of three analytical replicates.

**THF clathrate chamber design**. A cooling chamber was engineered with copper coils circulating ethylene glycol around a 50-mL glass beaker (**Fig. S2**). The base of the beaker was replaced with an acrylic disc (McMaster-Carr, 1221T63) with a side port for solution injection and extraction. The beaker was sealed with a rubber stopper with holes for a glass Pasteur pipette and a thermocouple connected to a Data Acquisition Unit (Keysight Technologies, GPIB, RS232).

**Experimental design.** 100% THF clathrate was synthesized by combining 19.1 wt% THF (Alfa Aesar, AA41820-AK) in deionized water^25^. All proteins were suspended in phosphate buffered saline (PBS, pH 7.53) solution. CbpAs were synthesized and purified within three days of each experiment to prevent protein decomposition. Cytochrome *c* (Sigma-Aldrich, C2506-50MG), which had previously been shown not to bind clathrate^26, 27^, and eGFP (cleaved from CbpA) were used as negative controls. Type I AFP (A/F Protein Inc., AFP Type I) was used as a positive control, as previously shown^26^. All proteins were tested at 100 µg mL^-1^ with few exceptions when protein concentration was not feasible.

**THF clathrate synthesis.** To begin the experiment, the chamber was cooled until solution temperature was stable between -2.5ºC and -4ºC. Treatment solution was injected through the side port. THF clathrate was nucleated by briefly inserting a dry ice-cooled copper wire through the glass Pasteur pipette and into the solution, 1 cm above the end of the pipette, as in previous studies^28, 29^. Crystal growth was imaged through a cutout in the cooling chamber using Nikon D7200 camera (Lens: AF-S DX VR Nikkor 18-55mm f/3.5-5.6G II) in an insulated box. A photograph was taken once every minute. When the crystal grew to the desired size (at least 1-2 cm; 2-4 hrs), the non-crystallized solution was extracted through the side port, revealing the THF clathrate crystal(s).

**Protein concentration in THF clathrate: non-crystallized solution.** THF clathrate melt and non-crystallized solution were weighed, THF was evaporated off in an anoxic gas stream, and samples were stored at 4ºC until spectrophotometric analysis. All samples were measured at 280 nm. eGFP-containing treatments were also measured at 491 nm. Cytochrome c was also measured at 550 nm. Using Beer’s law, we calculated the molar protein concentration from the absorbance value and the extinction coefficient (calculated using ExPasy^20^). Protein concentration in mg mL^-1^ was calculated by multiplying the molar concentration by the molecular weight of the protein. Protein concentration of the crystal melt was divided by protein concentration of the non-crystallized solution to obtain the ratio of protein that preferentially bound to the clathrate (**Fig. S3**).

**Clathrate freezing temperature and composition**. We note that although the freezing point of THF clathrate is +4.4ºC, we were unable to form solid above -2.5ºC at 19.1 wt% THF. In negative control experiments, crystals formed the same octahedral structure as previously synthesized THF clathrate crystals^26, 28, 29^ and are not similar to the most common hexagonal structure of water ice. We were unable to synthesize a crystal in the presence of water alone.

**Table S1. Accession numbers for Cbp proteins.**

| **Cbp** | **2** | **3** | **5** | **6** | **8** |
| --- | --- | --- | --- | --- | --- |
| **A** | ES708_13645 | GAI68542 | GAI60354 | ES708_30512 | ES707_98757 |
| **B** | ES708_13648 | GAI68555 | GAI60346 |  |  |
| **C** | ES708_13647 | GAI68551 | GAI60349 |  |  |
| **D** | ES708_13646 | GAI68546 | GAI60350 |  |  |
| **A'/"** |  | GAI68544 | GAI60352 |  |  |
| **Contig** | E10H5_C1877 | S12H4_C00034 | S12H4_C00157 | E10H5_C8338 | E5H5_C32933 |
| **Genome accession** | JABUBQ  010001866.1 | BARW  01000034.1 | BARW  01000157.1 | JABUBQ  010008252.1 | JABUBP  010032817.1 |

**Table S2. Amino acid frequency in CbpAs.**

|  | CbpA_2_ | | CbpA_3_ | | CbpA_5_ | | CbpA_6_ | | CbpA_8_ | |
| --- | --- | --- | --- | --- | --- | --- | --- | --- | --- | --- |
| Amino Acid | # | % | # | % | # | % | # | % | # | % |
| Ala (A) | 202 | 34.0 | 156 | 34.1 | 140 | 34.6 | 153 | 32.8 | 104 | 42.4 |
| Arg (R) | 22 | 3.7 | 27 | 5.9 | 27 | 6.7 | 30 | 6.4 | 18 | 7.3 |
| Asn (N) | 8 | 1.3 | 6 | 1.3 | 5 | 1.2 | 8 | 1.7 | 2 | 0.8 |
| Asp (D) | 15 | 2.5 | 6 | 1.3 | 5 | 1.2 | 8 | 1.7 | 0 | 0.0 |
| Cys (C) | 0 | 0.0 | 0 | 0.0 | 0 | 0.0 | 0 | 0.0 | 0 | 0.0 |
| Gln (Q) | 47 | 7.9 | 34 | 7.4 | 26 | 6.4 | 27 | 5.8 | 16 | 6.5 |
| Glu (E) | 72 | 12.1 | 56 | 12.2 | 52 | 12.8 | 61 | 13.1 | 36 | 14.7 |
| Gly (G) | 24 | 4.0 | 18 | 3.9 | 13 | 3.2 | 14 | 3.0 | 4 | 1.6 |
| His (H) | 4 | 0.7 | 3 | 0.7 | 2 | 0.5 | 2 | 0.4 | 1 | 0.4 |
| Ile (I) | 23 | 3.9 | 24 | 5.2 | 24 | 5.9 | 25 | 5.4 | 8 | 3.3 |
| Leu (L) | 24 | 4.0 | 17 | 3.7 | 13 | 3.2 | 21 | 4.5 | 3 | 1.2 |
| Lys (K) | 17 | 2.9 | 11 | 2.4 | 9 | 2.2 | 12 | 2.6 | 6 | 2.4 |
| Met (M) | 16 | 2.7 | 7 | 1.5 | 6 | 1.5 | 9 | 1.9 | 4 | 1.6 |
| Phe (F) | 4 | 0.7 | 5 | 1.1 | 2 | 0.5 | 2 | 0.4 | 0 | 0.0 |
| Pro (P) | 25 | 4.2 | 16 | 3.5 | 17 | 4.2 | 26 | 5.6 | 11 | 4.5 |
| Ser (S) | 9 | 1.5 | 15 | 3.3 | 12 | 3.0 | 12 | 2.6 | 1 | 0.4 |
| Thr (T) | 31 | 5.2 | 21 | 4.6 | 17 | 4.2 | 17 | 3.6 | 8 | 3.3 |
| Trp (W) | 16 | 2.7 | 6 | 1.3 | 7 | 1.7 | 11 | 2.4 | 5 | 2.0 |
| Tyr (Y) | 9 | 1.5 | 10 | 2.2 | 14 | 3.5 | 14 | 3.0 | 10 | 4.1 |
| Val (V) | 26 | 4.4 | 20 | 4.4 | 14 | 3.5 | 15 | 3.2 | 8 | 3.3 |
| Total | 594 | 100 | 458 | 100 | 405 | 100 | 467 | 100 | 245 | 100 |

\# indicates number of indicated amino acid type. % indicates percentage of amino acid type in overall CbpA.

**Table S3. *In silico* characterization of Cbp_2_ gene cluster.**

|  | **BLAST** | **HMMER** | **InterProt** | **Phyre2** |
| --- | --- | --- | --- | --- |
| **CbpA_2_** | NA | NA | NA | 30% confidence  5% coverage  membrane protein  PDB  5XW6-B |
| **CbpB_2_** | E = 3e^-57^  80% ID  60% coverage  TET42886  *Dehalococcoidia bacterium* | E = 2.4e^-54^  A0A523UK45_9CHLR  *Dehalococcoidia bacterium* | cysteine peptidase superfamily | 96% confidence  85% coverage  cysteine peptidase  PDB match 3KW0-D |
| **CbpC_2_** | E = 5e^0^  52% ID  37% coverage  OIW17195  *Lupinus angustifolius* | E = 8.9e^-5^, A0A2G9LZ84_9ARCH from  *Candidatus Woesearchaeota archaeon* | NA | 21% confidence  44% coverage  hydrolase  PDB  2QT7-B |
| **CbpD_2_** | E = 4e^-45^  42% ID  96% coverage  NIM45686 *Nitrososphaeria archaeon* | E = 8.7e^-27^, A0A497RS02_9EURY from  *Archaeoglobales archaeon* | NA | 66% confidence  18% coverage  hydrolase  PDB  1QDU-I |

**Table S4. *In silico* characterization of Cbp_3_ gene cluster.**

|  | **BLAST** | **HMMER** | **InterProt** | **Phyre2** |
| --- | --- | --- | --- | --- |
| **CbpA_3_** | NA | E = 4e^-11^  A0A133UGC4_9EURY  *Candidate division MSBL1 archaeon SCGC-AAA259E22* | NA | 34% confidence  32% coverage  LRR repeat  PDB match 3OJA-B |
| **CbpB_3_** | E = 3e^-59^  80% ID  62% coverage  TET42886 *Dehalococcoidia bacterium* | E = 8e^-56^  A0A523UK45_9CHLR  *Dehalococcoidia bacterium* | cysteine peptidase superfamily | 97% confidence  86% coverage  cysteine peptidase  PDB match 3KW0-D |
| **CbpC_3_** | E = 2e^-4^  34% ID  83% coverage  PIN71858  *Candidatus Woesearchaeota* archaeon | E = 6.8e^-5^  A0A2G9LZ84_9ARCH  *Candidatus Woesearchaeota* archaeon | NA | 19% confidence  21% coverage  transport protein  PDB match 5U1D-X |
| **CbpD_3_** | E = 2e^-43^  45% ID  89% coverage  NIM45686 *Nitrososphaeria* archaeon | E = 5.8e^-26^  A0A497RS02_9EURY *Archaeoglobales* archaeon | NA | 64% confidence  19% coverage  hydrolase  PDB match 1QDU-I |
| **CbpA_3’_** | E = 2e^-18^  39% ID  81% coverage  HHD66923  *Chloroflexi bacterium* | E = 7.2e^-59^  A0A133VL11_9EURY  *Candidate division MSBL1 archaeon SCGC-AAA382K21*, contains CARDB domain | CARDB Ig-like fold | 100% confidence  72% coverage  hydrolase  PDB match 5JP0-A |

**Table S5. Calculated biophysical parameters for CbpAs.**

|  | **eGFP-CbpA fusion**  **molecular weight (kDa)** | **CbpA molecular weight (kDa)** | **Extinction**  **coefficient (M^-1^cm^-1^)** |
| --- | --- | --- | --- |
| **CbpA_2_** | 91.2 | 61.3 | 101,410 |
| **CbpA_3_** | 77.1 | 47.2 | 47,900 |
| **CbpA_5_** | 72.2 | 42.3 | 59,360 |
| **CbpA_6_** | 79.1 | 49.2 | 81,360 |

**Table S6.** **Key biophysical properties of CbpAs.**

|  | **Target Freeze score** | **Secondary structure** | **Melting temperature (°C)** |
| --- | --- | --- | --- |
| **CbpA_2_** | 0.924 | alpha helix | 33.5 |
| **CbpA_3_** | 0.912 | alpha helix | 38.0 |
| **CbpA_5_** | 0.907 | alpha helix | 33.3 |
| **CbpA_6_** | 0.905 | alpha helix | 36.0 |
| **CbpA_8_** | 0.834 | NA | NA |

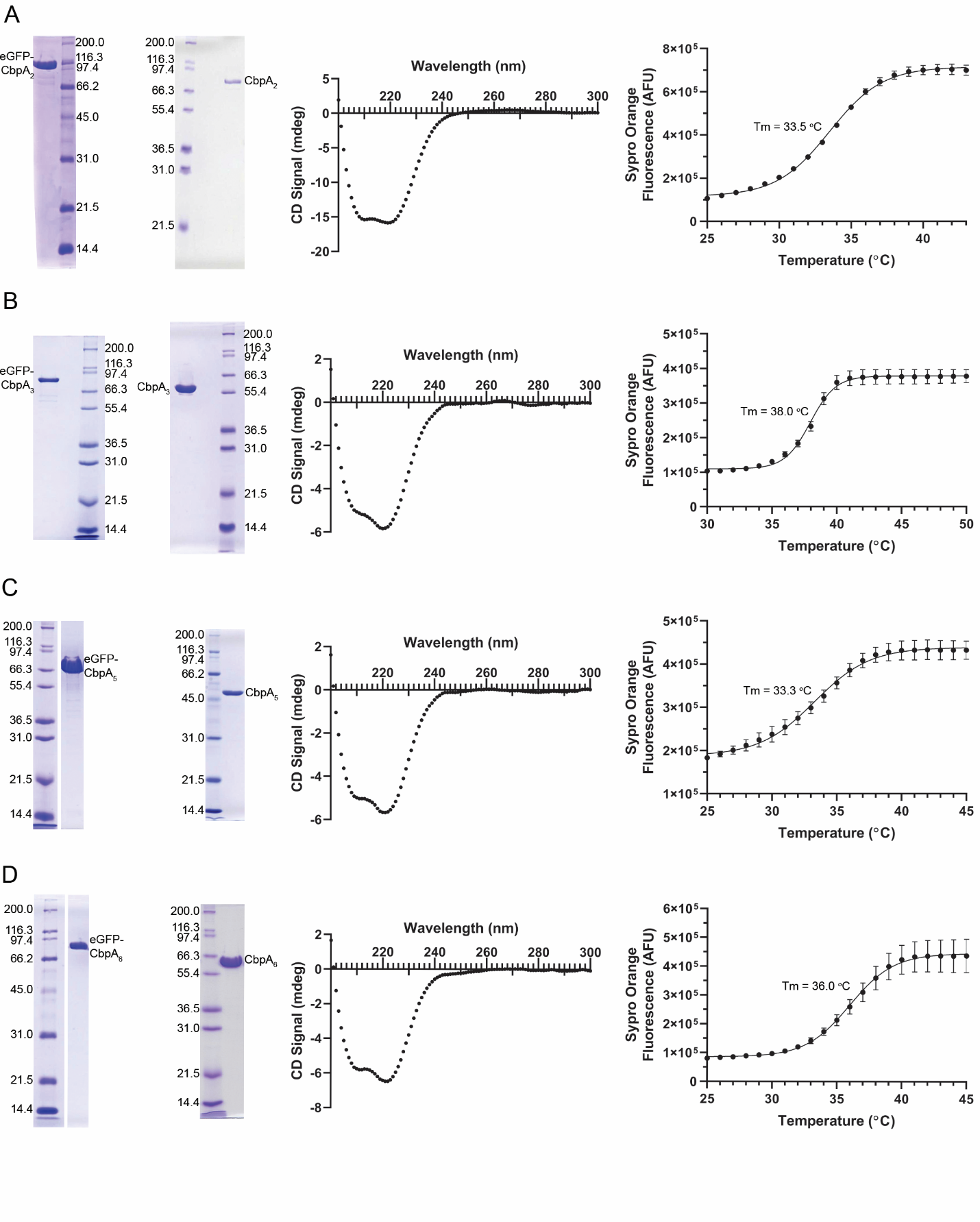

**Figure S1. Biophysical characterization of clathrate binding proteins**. Characterization of CbpA_2_ (A), CbpA_3_ (B), CbpA_5_ (C), and CbpA_6_ (D). *Left:* SDS-PAGE analysis of purified eGFP-CbpA and CbpAs. *Middle:* CD spectra showing that all CbpAs described herein have alpha-helical secondary structure elements. *Right:* DSF thermal melting curves with melting temperature (Tm) indicated for each Cbp_A_ protein. All T_m_ values are < 40°C for CbpAs in 10 mM HEPES at pH 7.2 with 0.2 M NaCl.

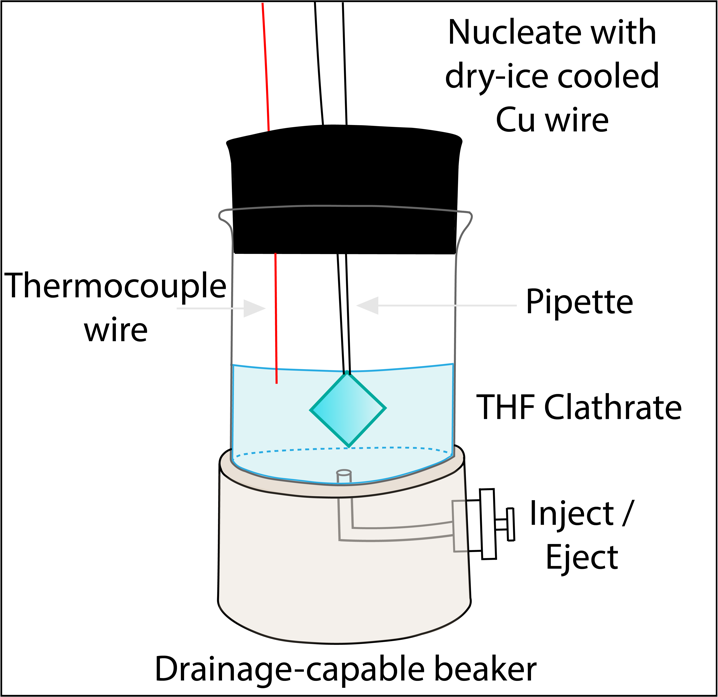

**Figure S2. A glass beaker with an acrylic base was used to synthesize THF clathrate in the presence of CbpAs.** 19.1wt% THF solution containing the treatment was prepared and injected through the side port. The solution was cooled and THF clathrate was nucleated with a dry ice-cooled Cu wire down the pipette about 1 cm above the end of the pipette. The non-crystallized solution was ejected through the side port, revealing the clathrate crystal(s). The clathrate was imaged and the protein was quantified using spectrophotometric analysis.

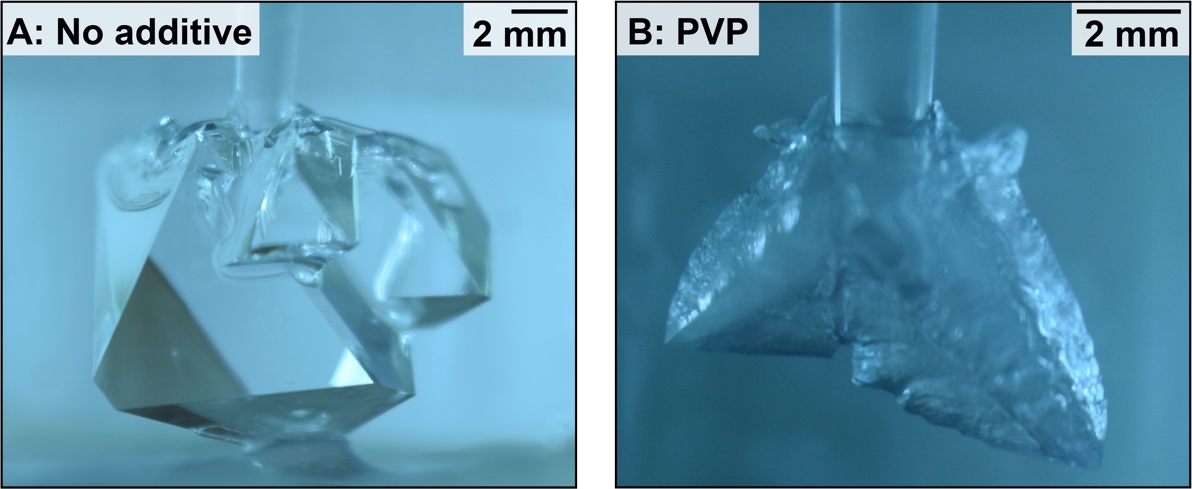

**Figure S3. Control treatments for THF clathrate** conducted with no additive (A), yielding regular, twinned clathrate crystals, and (B) with polyvinylpyrollidone (PVP; 1 mg mL^-1^), yielding a skeletal clathrate crystal, as shown in previous studies^26^.

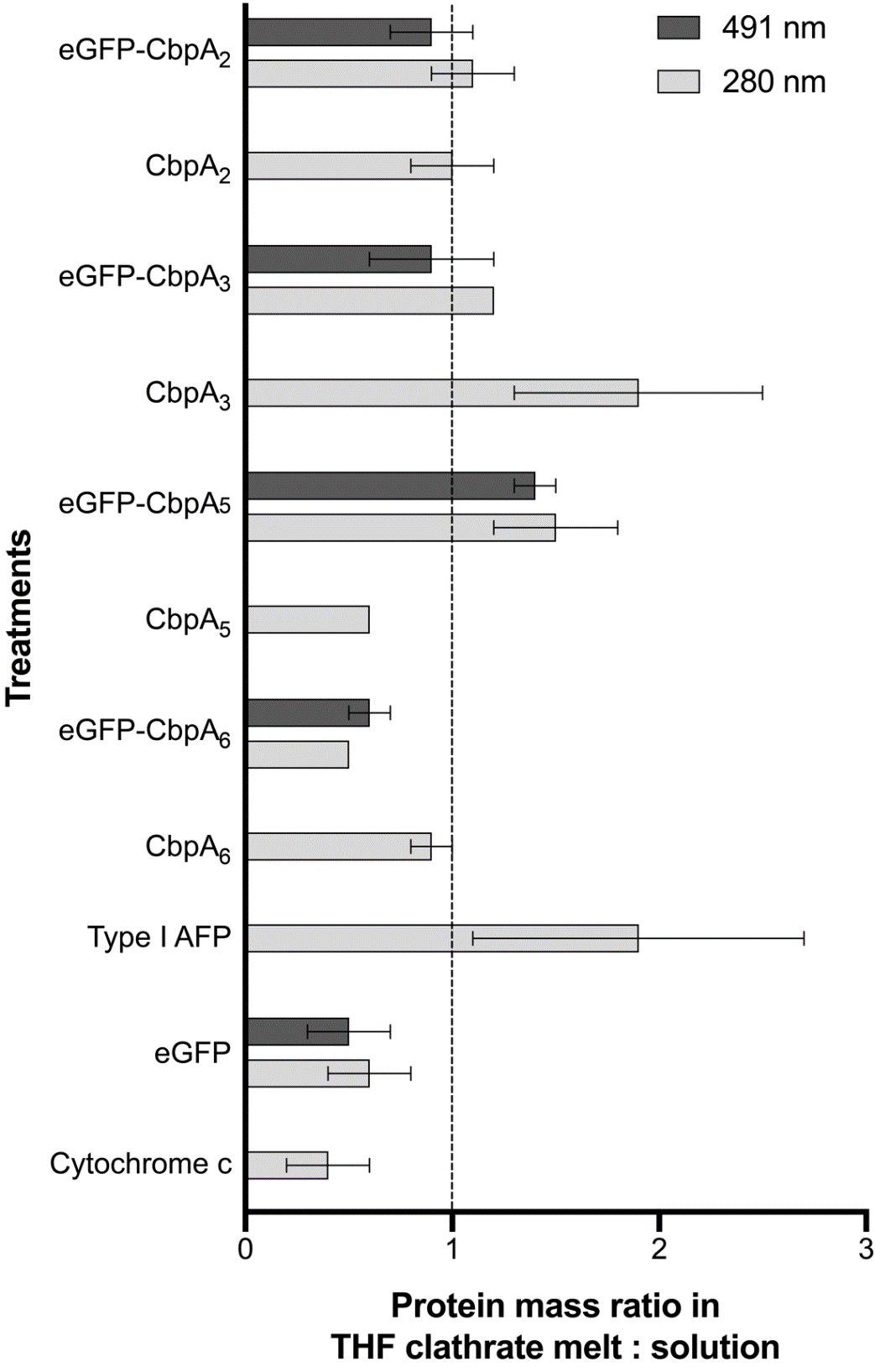

**Figure S4. CbpAs preferentially bind to THF clathrate*.*** The average protein mass ratio in THF clathrate crystal vs. the non-crystallized solution calculated from spectrophotometric analysis is shown. Negative controls are cytochrome c and eGFP. Type I AFP is the positive control. CbpAs and eGFP-CbpAs were analyzed. All treatments were analyzed via 280 nm. eGFP-CbpAs and eGFP were also analyzed at 491 nm. Values greater than 1 indicate higher protein concentration in the crystal than in the non-crystallized solution.

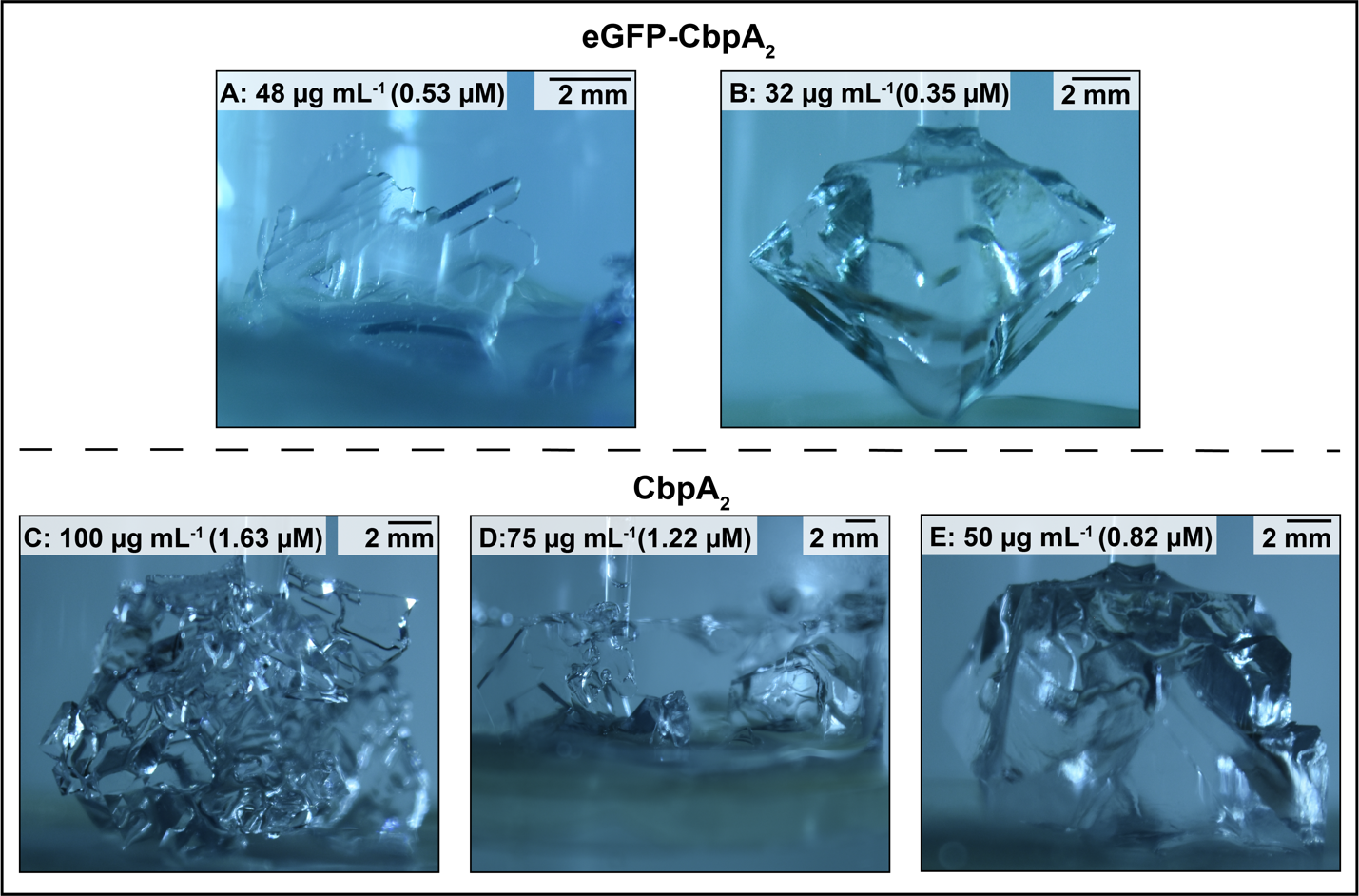

**Figure S5. Increasing CBP concentration induces greater change in clathrate morphology.** eGFP-CbpA_2_ (top) and CbpA_2_ (bottom) show more morphologic alteration of THF clathrate at higher concentrations. THF clathrate exhibited flat plate-like morphology with jagged edges in an eGFP-CbpA_2_ trial at 48 µg mL^-1^ (0.53 µM, A); however, at 32 µg mL^-1^ (0.35 µM, B), THF clathrate formed a single crystal with surface deformities, such as kinking. For CbpA_2_ treatments, we completed two trials at 100 µg mL^-1^ (1.63 µM, C), one trial at 75 µg mL^-1^ (1.22 µM, D), and one trial at 50 µg mL^-1^ (0.82 µM, E). The trials with 100 and 75 µg mL^-1^ yielded polycrystalline, plate-like morphology, whereas the 50 µg mL^-1^ treatment yielded a single THF clathrate crystal with surficial deformities.

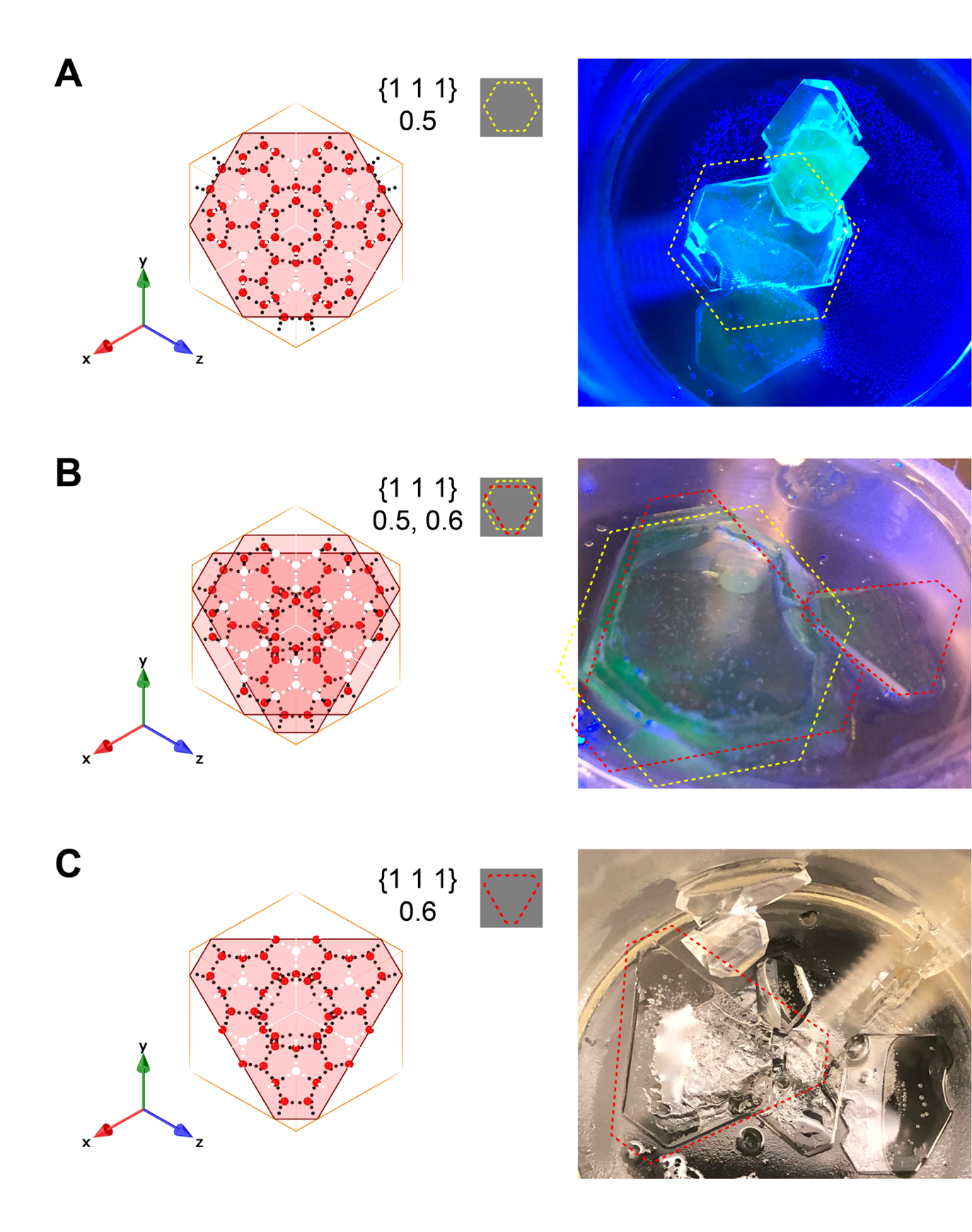

**Figure S6. CBPs induce platelike clathrate growth along the [1 1 1] plane.** eGFP-CbpA_6_ (A), eGFP-CbpA_5_ (B), and CbpA_6_ (C) promoted clathrate growth in hexagonal sheets, similar to the shape of a regular octahedral THF clathrate crystal at depths 0.5 and 0.6 observed in Crystal Maker v10.5.2. The left column depicts the shape of the [1 1 1] face in the 0.5 depth (A), 0.5 and 0.6 depths (B), and 0.6 depth (C), with red dots representing hydrogen-bonded waters that form cages around THF molecules. The small grey box contains the outline of the 0.5 (yellow dashed) and 0.6 (red-dashed) depth hexagons, which are used to outline the crystals in the right column.

**References.**

1.	Glass, J. B.; Ranjan, P.; Kretz, C. B.; Nunn, B. L.; Johnson, A. M.; McManus, J.; Stewart, F. J., Adaptations of Atribacteria to life in methane hydrates: hot traits for cold life. *Environ. Microbiol.*

8.	Foundation, P. S. *Python Language Reference*, 3.7.

9.	Doxey, A. C.; Yaish, M. W.; Griffith, M.; McConkey, B. J., Ordered surface carbons distinguish antifreeze proteins and their ice-binding regions. *Nat Biotechnol* **2006,** *24* (7), 852-5.

10.	He, X.; Han, K.; Hu, J.; Yan, H.; Yang, J. Y.; Shen, H. B.; Yu, D. J., TargetFreeze: Identifying Antifreeze Proteins via a Combination of Weights using Sequence Evolutionary Information and Pseudo Amino Acid Composition. *J Membr Biol* **2015,** *248* (6), 1005-14.

11.	Consortium, T. U., UniProt: a worldwide hub of protein knowledge. *Nucleic Acids Research* **2018,** *47* (D1), D506-D515.

12.	Mitchell, A. L.; Attwood, T. K.; Babbitt, P. C.; Blum, M.; Bork, P.; Bridge, A.; Brown, S. D.; Chang, H. Y.; El-Gebali, S.; Fraser, M. I.; Gough, J.; Haft, D. R.; Huang, H.; Letunic, I.; Lopez, R.; Luciani, A.; Madeira, F.; Marchler-Bauer, A.; Mi, H.; Natale, D. A.; Necci, M.; Nuka, G.; Orengo, C.; Pandurangan, A. P.; Paysan-Lafosse, T.; Pesseat, S.; Potter, S. C.; Qureshi, M. A.; Rawlings, N. D.; Redaschi, N.; Richardson, L. J.; Rivoire, C.; Salazar, G. A.; Sangrador-Vegas, A.; Sigrist, C. J. A.; Sillitoe, I.; Sutton, G. G.; Thanki, N.; Thomas, P. D.; Tosatto, S. C. E.; Yong, S. Y.; Finn, R. D., InterPro in 2019: improving coverage, classification and access to protein sequence annotations. *Nucleic Acids Res* **2019,** *47* (D1), D351-D360.

13.	Potter, S. C.; Luciani, A.; Eddy, S. R.; Park, Y.; Lopez, R.; Finn, R. D., HMMER web server: 2018 update. *Nucleic Acids Res* **2018,** *46* (W1), W200-W204.

20.	Walker, J. M., *The proteomics protocols handbook*. Humana Press: Totowa, N.J., 2005; p xviii, 988 p.

26.	Gordienko, R.; Ohno, H.; Singh, V. K.; Jia, Z.; Ripmeester, J. A.; Walker, V. K., Towards a green hydrate inhibitor: imaging antifreeze proteins on clathrates. *PLoS One* **2010,** *5* (2), e8953.

27.	Zeng, H.; Wilson, L. D.; Walker, V. K.; Ripmeester, J. A., Effect of antifreeze proteins on the nucleation, growth, and memory effect during tetrahydrofuran clathrate hydrate formation. *J. Am. Chem. Soc.* **2006,** *128*, 2844-2850.

28.	Makogon, T. Y.; Larsen, R.; Knight, C. A.; Sloan, E. D., Melt growth of tetrahydrofuran clathrate hydrate and its inhibition: method and first results. *Journal of Crystal Growth* **1997,** *179*, 258-262.

29.	Zeng, H.; Wilson, L. D.; Walker, V. K.; Ripmeester, J. A., The inhibition of tetrahydrofuran clathrate-hydrate formation with antifreeze protein. *Canadian Journal of Physics* **2003,** *81* (1-2), 17-24.
